# Sexual Conflict through Mother’s Curse and Father’s Curse

**DOI:** 10.1101/345611

**Authors:** J. Arvid Ågren, Manisha Munasinghe, Andrew G. Clark

**Affiliations:** Department of Molecular Biology and Genetics, Cornell University, Ithaca, NY, 14583, USA; Department of Biological Statistics and Computational Biology, Cornell University, Ithaca, NY, 14853, USA

**Author notes:** These authors contributed equally (J. Arvid Ågren, Manisha Munasinghe).

**Keywords:** Genomic conflict, Sex chromosomes, Mitochondria, Sexual antagonism

## Abstract

In contrast with autosomes, lineages of sex chromosomes reside for different amounts of time in males and females, and this transmission asymmetry makes them hotspots for sexual conflict. Similarly, the maternal inheritance of the mitochondrial genome (mtDNA) means that mutations that are beneficial in females can spread in a population even if they are deleterious in males, a form of sexual conflict known as Mother’s Curse. While both Mother’s Curse and sex chromosome induced sexual conflict have been well studied on their own, the interaction between mitochondrial genes and genes on sex chromosomes is poorly understood. Here, we use analytical models and computer simulations to perform a comprehensive examination of how transmission asymmetries of nuclear, mitochondrial, and sex chromosome-linked genes may both cause and resolve sexual conflicts. For example, the accumulation of male-biased Mother’s Curse mtDNA mutations will lead to selection in males for compensatory nuclear modifier loci that alleviate the effect. We show how the Y chromosome, being strictly paternally transmitted provides a particularly safe harbor for such modifiers. This analytical framework also allows us to discover a novel kind of sexual conflict, by which Y chromosome-autosome epistasis may result in the spread of male beneficial but female deleterious mutations in a population. We christen this phenomenon Father’s Curse. Extending this analytical framework to ZW sex chromosome systems, where males are the heterogametic sex, we also show how W-autosome epistasis can lead to a novel kind of nuclear Mother’s Curse. Overall, this study provides a comprehensive framework to understand how genetic transmission asymmetries may both cause and resolve sexual conflicts.

## Introduction

Males and females differ in a wide variety of phenotypic traits, including reproduction, behavior, and morphology (Darwin, 1871). Because the sexes largely share the same genome, an allele that moves a trait toward the optimal value for one sex, may have negative effect on the fitness of the other sex, a phenomenon now known as intra-locus sexual conflict (Arnqvist and Rowe, 2005; Parker and Others, 1979; Pennell and Morrow, 2013; Trivers, 1972). Phenotypic assays in *Drosophila* (Chippindale et al., 2001) and water striders (Arnqvist and Rowe, 2002) provided early direct empirical support for the occurrence and dynamic nature of such conflicts. More recently, rapid advancements in sequencing technology means that these questions can now be tackled molecularly, leading to a revived interest in the topic (reviewed in e.g. (Kasimatis et al., 2017; Mank, 2017; Wright and Mank, 2013)). Several studies have examined the genomics of sexual conflict (Cheng and Kirkpatrick, 2016; Innocenti and Morrow, 2010; Wright et al., 2018), however the exact nature of these signatures are still poorly understood.

The genetics of sexual conflicts is further complicated by the parts of the genome not equally shared by males and females. In contrast with autosomes, lineages of sex chromosomes spend different amounts of time in males and females and this transmission asymmetry makes them hotspots for sexual conflict (Jaenike, 2001; Mank et al., 2014; van Doorn and Kirkpatrick, 2007). For example, in species with a XY sex chromosome system and a 50:50 sex ratio, a given X chromosome spends 2/3 of its time in females. This has led to the prediction that the X chromosome should become “feminized” or “demasculinized”, which may occur through inter-chromosomal translocations, gene duplications, or alterations in sex-specific gene expression (Connallon and Clark, 2011; Gallach and Betrán, 2011). Consistent with this, an overrepresentation of female ovary-specific, and an under-representation of male testis-specific genes on the X has been reported in several *Drosophila* species (Allen et al., 2013; Meisel et al., 2012; Sturgill et al., 2007), though not in, for example, humans or mice (Lercher et al., 2003; Yang et al., 2006)). Analogously, in an ZW sex chromosome system, where females are the heterogametic sex, we would then expect to see a masculinization of the Z chromosome (Rice, 1984). In line with this prediction, a series of gene expression studies of the Z have reported a trend of male-biased expression (Kaiser and Ellegren, 2006; Mank and Ellegren, 2009; Storchová and Divina, 2006).

By similar logic, the maternal inheritance of the mitochondrial genome (mtDNA) means that mutations that are beneficial in females can spread in a population even if they are deleterious in males (Charlesworth and Charlesworth, 1978; Frank, 1989; Frank and Hurst, 1996; Gemmell et al., 2004; Lewis, 1941; Vaught and Dowling, 2018). This has been particularly well studied in flowering plants, where mitochondrial mutations that prevent pollen production in otherwise hermaphroditic plants are widespread, a phenomenon known as cytoplasmic male sterility (Budar et al., 2003; Case et al., 2016; Lewis, 1941). Selection on male function, so-called nuclear restorers of fertility, has then led to a co-evolutionary arms race, which may result in reproductive isolation of diverged populations (reviewed in (Ågren 2013; Case et al., 2016; Crespi and Nosil, 2013)). Similar examples from animal systems were long missing. However, during the last few years, a series of studies have provided evidence of the occurrence of Mother’s Curse, as this form of sexual conflict is typically referred to in animals (Gemmell et al., 2004). Experimental work in *Drosophila melanogaster* has demonstrated that males are more sensitive to mitochondrial genetic variation than females, which has been interpreted as evidence of Mother’s Curse (Camus et al., 2012). Furthermore, in a technical *tour de force* with *D. melanogaster*,Patel et al. (2016) identified COII^G177S^, a mitochondrial hypomorph of cytochrome oxidase II, as a male-harming locus causing a reduction in male fertility through a disruption of sperm development, but without any negative effects in females. It is worth noting that the male sterile effect of COII^G177S^ is dependent on the nuclear background. Such epistasis is in line with earlier suggestions that the accumulation of male-harming mitochondrial mutations should, just like in the plant cytoplasmic male sterility example, result in nuclear-encoded restorers of male fitness (Dowling et al., 2008; Rand et al., 2004). Finally, Mother’s Curse has also recently been unequivocally demonstrated in humans, in the example of the mitochondrial mutation resulting in Leber’s hereditary optic neuropathy, the degradation of retinal ganglion cells, resulting in loss of vision, which predominantly affects males (Milot et al., 2017).

Thus, mitochondrial, X and Y, and Z and W, genes have all been shown to be involved in sexual conflicts. However, while these examples have received both theoretical and empirical attention on their own, we lack a clear picture of how mitochondrial, nuclear, and sex linked genes may interact in sexually antagonistic ways. Here, we develop a comprehensive theoretical framework to examine the role of genetic transmission asymmetries in sexual conflicts. We consider the sexually antagonistic consequences of mitochondrial-autosome, mitochondrial-sex chromosome, and autosome-sex chromosome interactions in XY and ZW sex chromosome systems. These models show how the fate of a nuclear allele restoring male fitness in the face of Mother’s Curse depends on whether it is located on autosomes, X, Y, or Z. Our models also allow us to discover a novel kind of sexual conflict, by which Y chromosome-autosome, or Y-X, epistasis may result in the spread of male beneficial but female deleterious mutations in a population. We name this phenomenon Father’s Curse. Analogously, we find that W-autosome, or W-Z interactions may also result in a nuclear version of the Mother’s Curse. Taken together, our results extend previous theoretical work on how genetic transmission asymmetries may lead to sexually antagonistic selection, but also on occasion reduce sexual conflict.

### Modeling framework

We consider nine two-locus two-allele models to investigate the fitness consequences for males and females of epistatic interactions between mitochondria and autosomes, mitochondria and sex chromosomes, and between sex chromosomes and autosomes (Fig. 1). We perform this analysis in both XY and ZW sex chromosomes systems. For each of the nine models below, we first discuss previous theoretical work and the available empirical data, and then outline our model and results. Genotypes and fitnesses for males and females in all models are summarized in Table 1.

**Figure 1.**
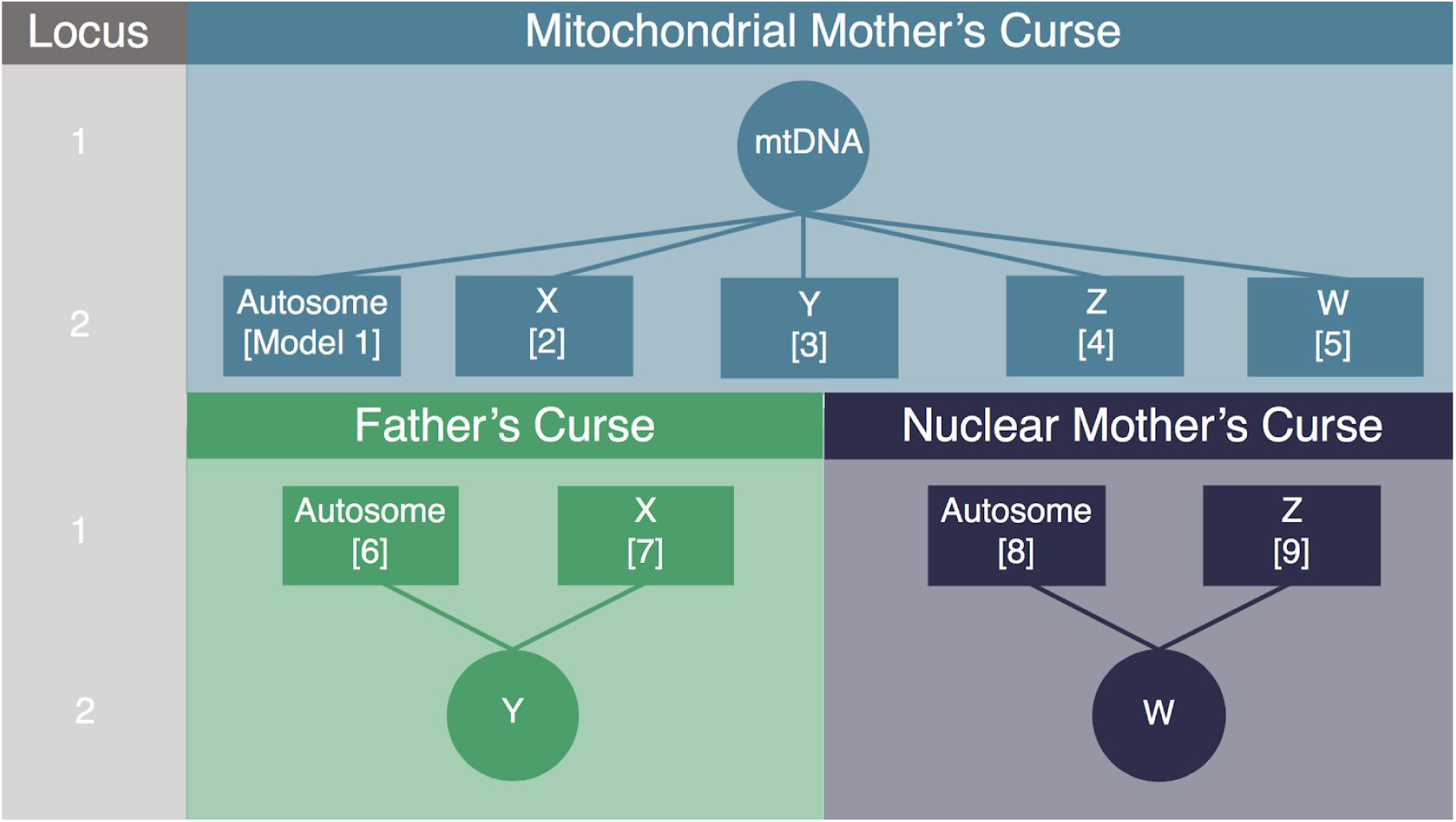
Graphical representation of the 9 two-locus two-allele models considered. Models 1-5 capture the dynamics of a mitochondrial Mother’s Curse mutation and a nuclear restorer located on an autosome or sex chromosome respectively. In each model, Locus 1 is the primary sex-asymmetric locus, and Locus 2 harbors alleles that act to restore the fitness of the disadvantaged sex, except Model 5 where it further contributes to the advantaged sex. In Models 6-9, we model how selection on a mutation on the uniparentally inherited sex chromosome (Y, Models 6-7; W, Models 8-9) can lead to the spread of a mutation that is beneficial in the heterogametic sex but deleterious in the homogametic sex, Father’s Curse and Nuclear Mother’s Curse respectively.

**Table 1.**
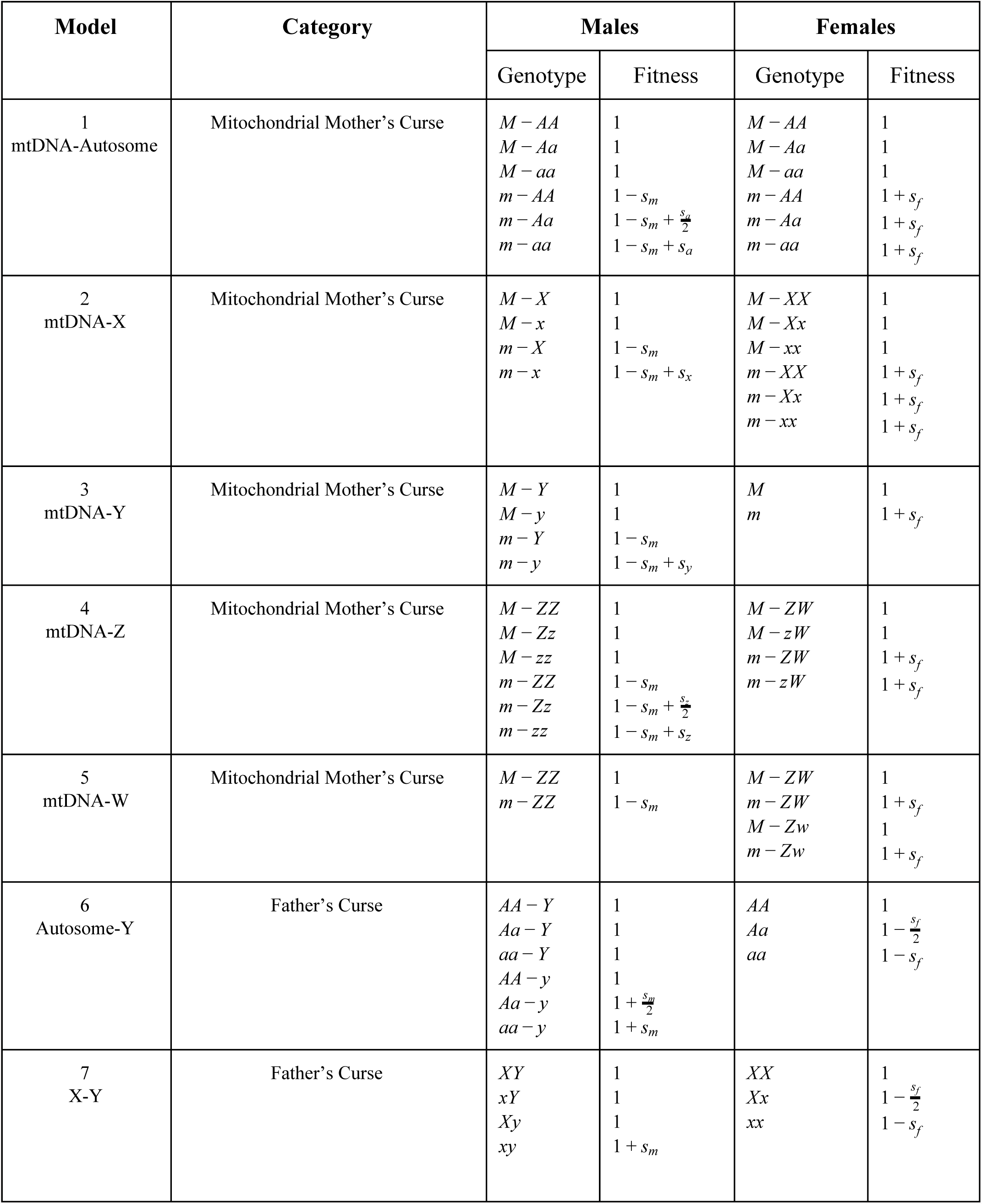
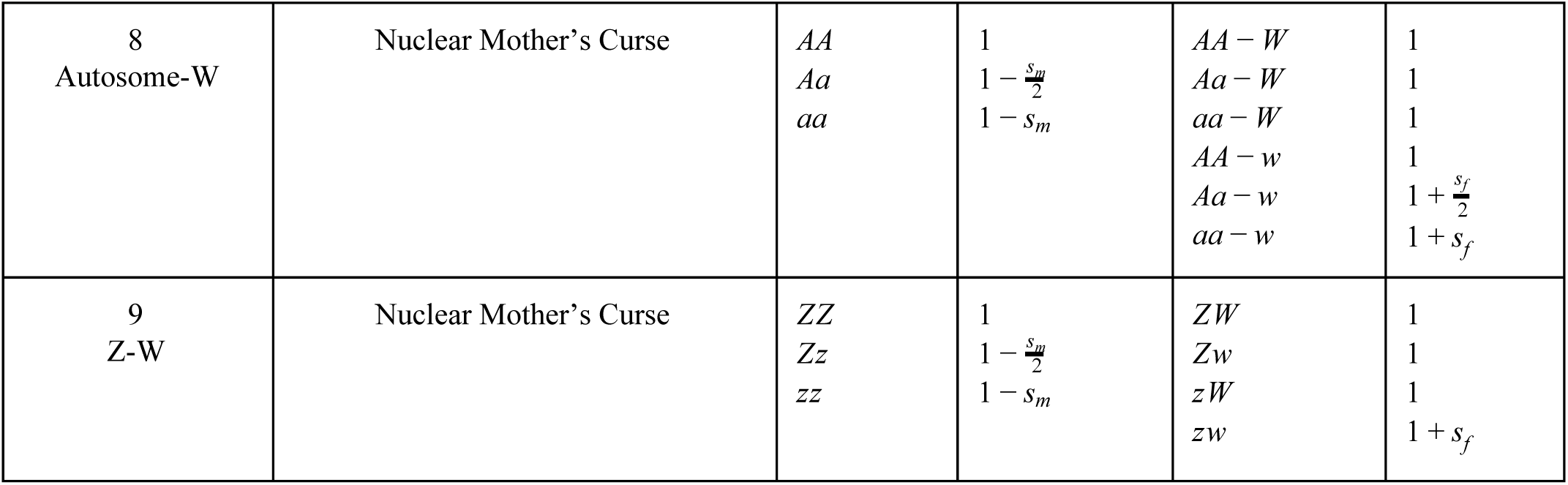
Genotypes and fitnesses in Models 1-9

| Model | Category | Males |  | Females |  |
| --- | --- | --- | --- | --- | --- |
|  |  | Genotype | Fitness | Genotype | Fitness |
| 1<br>mtDNA-Autosome | Mitochondrial Mother's Curse | $M - AA$<br>$M - Aa$<br>$M - aa$<br>$m - AA$<br>$m - Aa$<br>$m - aa$ | 1<br>1<br>1<br>$1 - s_m$<br>$1 - s_m + \frac{s_a}{2}$<br>$1 - s_m + s_a$ | $M - AA$<br>$M - Aa$<br>$M - aa$<br>$m - AA$<br>$m - Aa$<br>$m - aa$ | 1<br>1<br>1<br>$1 + s_f$<br>$1 + s_f$<br>$1 + s_f$ |
| 2<br>mtDNA-X | Mitochondrial Mother's Curse | $M - X$<br>$M - x$<br>$m - X$<br>$m - x$ | 1<br>1<br>$1 - s_m$<br>$1 - s_m + s_x$ | $M - XX$<br>$M - Xx$<br>$M - xx$<br>$m - XX$<br>$m - Xx$<br>$m - xx$ | 1<br>1<br>1<br>$1 + s_f$<br>$1 + s_f$<br>$1 + s_f$ |
| 3<br>mtDNA-Y | Mitochondrial Mother's Curse | $M - Y$<br>$M - y$<br>$m - Y$<br>$m - y$ | 1<br>1<br>$1 - s_m$<br>$1 - s_m + s_y$ | $M$<br>$m$ | 1<br>$1 + s_f$ |
| 4<br>mtDNA-Z | Mitochondrial Mother's Curse | $M - ZZ$<br>$M - Zz$<br>$M - zz$<br>$m - ZZ$<br>$m - Zz$<br>$m - zz$ | 1<br>1<br>1<br>$1 - s_m$<br>$1 - s_m + \frac{s_z}{2}$<br>$1 - s_m + s_z$ | $M - ZW$<br>$M - zW$<br>$m - ZW$<br>$m - zW$ | 1<br>1<br>$1 + s_f$<br>$1 + s_f$ |
| 5<br>mtDNA-W | Mitochondrial Mother's Curse | $M - ZZ$<br>$m - ZZ$ | 1<br>$1 - s_m$ | $M - ZW$<br>$m - ZW$<br>$M - Zw$<br>$m - Zw$ | 1<br>$1 + s_f$<br>1<br>$1 + s_f$ |
| 6<br>Autosome-Y | Father's Curse | $AA - Y$<br>$Aa - Y$<br>$aa - Y$<br>$AA - y$<br>$Aa - y$<br>$aa - y$ | 1<br>1<br>1<br>1<br>$1 + \frac{s_m}{2}$<br>$1 + s_m$ | $AA$<br>$Aa$<br>$aa$ | 1<br>$1 - \frac{s_f}{2}$<br>$1 - s_f$ |
| 7<br>X-Y | Father's Curse | $XY$<br>$xY$<br>$Xy$<br>$xy$ | 1<br>1<br>1<br>$1 + s_m$ | $XX$<br>$Xx$<br>$xx$ | 1<br>$1 - \frac{s_f}{2}$<br>$1 - s_f$ |
| 8<br>Autosome-W | Nuclear Mother's Curse | $AA$<br>$Aa$<br>$aa$ | 1<br>$1 - \frac{s_m}{2}$<br>$1 - s_m$ | $AA - W$<br>$Aa - W$<br>$aa - W$<br>$AA - w$<br>$Aa - w$<br>$aa - w$ | 1<br>1<br>1<br>1<br>$1 + \frac{s_f}{2}$<br>$1 + s_f$ |
| 9<br>Z-W | Nuclear Mother's Curse | $ZZ$<br>$Zz$<br>$zz$ | 1<br>$1 - \frac{s_m}{2}$<br>$1 - s_m$ | $ZW$<br>$Zw$<br>$zW$<br>$zw$ | 1<br>1<br>1<br>$1 + s_f$ |

Most of these models feature a biallelic locus that generates the sexual conflict and another biallelic locus that acts to restore fitness to the disadvantaged sex. With the resulting four gametic types, the models typically have 3 degrees of freedom, and can be described by allele frequencies at the two loci, plus a linkage-disequilibrium-like term (Clark 1984; Asmussen et al. 1987). None of these models harbor a joint polymorphism at both loci, so our only interest is in stability of the fixation states, where the disequilibrium terms collapse to zero. For simplicity, we consider the two-dimension systems characterized by the allele frequencies at the two loci, and we determine local stability from the leading eigenvalue of the linearized recursion.

To further confirm the analytical results, we modeled selection using forward simulations that incorporated selection as a deterministic process with initial zygote frequencies, followed by selection which acts as weights on the frequencies of the zygotes, and then random mating and gamete formation following Mendelian rules. In each model, the simulations track all possible genotypes in each sex (which are denoted for each model in Table 1). Mitochondrial Mother’s Curse simulations (Models 1-4) track the frequency and time to fixation of the mutant alleles for a given parameter set *s_f_*, *s_m_*, *s_x_* of *s_z_* for an and arbitrary 5000 generations. Similarly, Father’s Curse and nuclear Mother’s Curse track the frequency and time to fixation of the mutant alleles for a given parameter set of *s_f_* and *s_m_* for 10,000 generations. All simulations were run in R v. 3.3.1, and scripts are in GitHub (https://github.com/mam737/ParentalCurseScripts).

## Models

### Mitochondrial Mother’s Curse

#### Model 1 Mitochondrial Mother’s Curse and Autosomal restoration

Early theoretical work on mito-autosome interactions was done in the 1970s (Charlesworth and Charlesworth, 1978; Charlesworth and Ganders, 1979) and 1980s (Arnold et al., 1988; Asmussen et al., 1987; Charlesworth, 1981; Clark, 1984). Models of cytoplasmic male sterility as an evolutionary route from hermaphroditism to dioecy, via gynodioecy (the presence of hermaphrodites and females) has also received extensive attention in flowering plants, and these models have since accumulated abundant empirical support (Charlesworth et al., 2010; Delph et al., 2007; Saur Jacobs and Wade, 2003). Furthermore, the importance of mito-nuclear compatibility in flowering plants is also revealed by the large number of cases of mito-nuclear induced reproductive isolation reported (Bomblies, 2010; Rieseberg and Blackman, 2010). More recently, several theoretical models of male fitness restoration in response to Mother’s Curse have been developed (Connallon et al., 2018; Unckless and Herren, 2009; Wade and Brandvain, 2009; Wade and Drown, 2016). This upswing in interest is partially due to an increased general appreciation of the importance of mitochondrial variation on fitness (Baris et al., 2017; Camus et al., 2012; Mossman et al., 2016), as well as direct test of the Mother’s Curse (Milot et al., 2017; Patel et al., 2016). Thus, the basic population dynamics of mito-autosome interactions are well worked out, so only for completeness do we lay out the model here.

There are two mitochondrial types (*M* and *m*) such that the ancestral type (*M*) has fitness 1 and the derived type (*m*) has an advantage in females and disadvantage in males. There is one autosomal locus that can serve as a restorer of fitness in males, so each sex has six cytogenotypes (*M-AA*, *M-Aa*, *M-aa*, *m-AA*, *m-Aa* and *m-aa*; Table 1). The fitnesses of all females with the *M* mitotype are 1 regardless of the autosomal locus, and those with the *m* mitotype have fitness 1+ *s_f_*. Male genotypes with the *M* mitotype also all have fitness 1, and those with the *m* mitochondria have fitnesses (in the order as above: 1, 1-*s_m_*+*s_a_*)2, and 1-*s_m_*+*s_a_*), where *s_m_* is the deleterious male effect of the new mitochondrial type and *s_a_* is the effect of the restorer (which is assumed to be additive). By normalizing frequencies within each sex, we are implicitly assuming that males can inseminate all reproductive females, and that a skewed sex ratio plays no role in the dynamics of either locus.

Whenever *s_f_* > 0, the *m* = 0 equilibrium is unstable, and the Mother’s Curse mitochondrial type increases in frequency all the way to fixation, regardless of the fitnesses of the male genotypes. The conditions for invasion of the compensatory *a* allele depend on the frequency of *m*, but once the *m* allele has gone to fixation, the condition for invasion is simply *sa* > *s_m_* (Fig. 2). Connallon et al. (2017) consider the male genetic load in a similar model, noting that the load depends on the arrival interval of both mitochondrial and autosomal mutations with appropriate fitness effects, as well as the distribution of fitness effects of those mutations. The model suggests that the interesting empirical question that remains is to better quantify these mutation rates and distributions of fitness effects.

**Figure 2.**
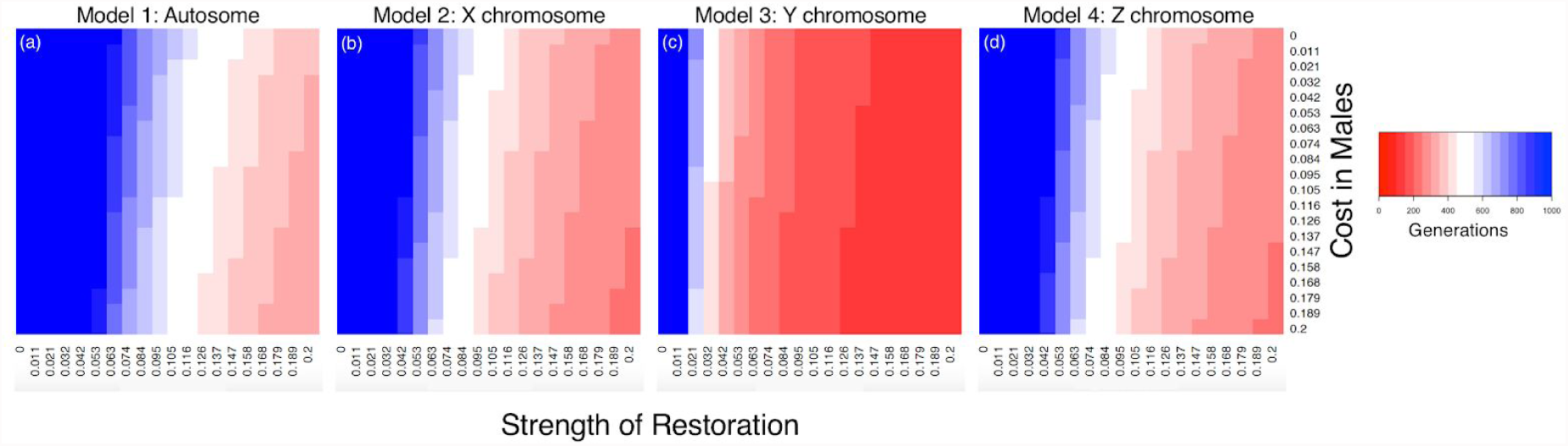
Fixation time for Mother’s Curse compensators. The rate of fixation for a compensator located on an autosome (a; Model 1), X chromosome (b; Model 2), Y chromosome (c; Model 3), or Z chromosome (d; Model 4) is contrasted for a given female benefit (*s_f_* = 0.105; simulations were ran for 19 different values evenly spaced between 0.02 and 0.4) is shown. Colors show the number of generations (up to 1000) required for nuclear compensator that restores male fitness to fix. The x-axis is the strength of restoration of the compensator (*s_a_*, *s_x_*, *s_y_*, and *s_z_* respectively).

Model 1 as stated has highly constrained fitnesses, and one might ask how the dynamics change when, for example, there is coadaptation between mitochondrial and autosomal loci, *i.e*. the *M-AA* and *m-aa* female genotypes both have the highest fitnesses. Clark (1984) showed that, in a one-sex model with autosomal and mitochondrial loci, regardless of the configuration of the six fitnesses, there is no admissible stable polymorphism for both loci.

#### Model 2 Mitochondrial Mother’s Curse and X-linked restoration

The co-evolution between mitochondrial and X-linked genes has been modelled by Rand et al. (2001), as well as by Wade and Drown (2016). These models highlight how mito-X epistasis can maintain mito-nuclear polymorphisms, and because the X spends 2/3 of its time in females it can exacerbate the spread of male-harming mitochondrial mutations. Rand et al. (2001) also used experiments in *Drosophila melanogaster* to provide direct empirical support for the importance of mito-X interactions. Further indirect evidence of mito-X interactions come from studies of the genomic distribution of nuclear genes with mitochondrial functions, which are assumed to have migrated from the mitochondrial to the nuclear genome. Early evidence suggested a general pattern by which such genes were under-represented on the X across multiple taxa (Drown et al., 2012). However, later work using phylogenetic independent methods demonstrated that this pattern was restricted to therian mammals and *Caenorhabditis elegans* (Dean et al., 2014). All other animal species studied show no chromosomal bias in their distribution. The same lack of biased distribution was also seen in a dioecious plant with sex chromosomes (Hough and Ågren et al. 2014).

Again, we have mitochondrial alleles (*M* and *m*) and now a pair of X chromosomal alleles (*X* and *x*). Females are of six genotypes (*M-XX*, *M-Xx*, *M-xx*, *m-XX*, *m-Xx*, and *m-xx*), and males are of four genotypes (*M-XY*, *M-xY*, *m-XY*, and *m-xY*). In males, the *m* mitochondrial allele comes with a fitness cost, but only with the ancestral X chromosome, and the *x* allele provides fitness compensation of *s_x._* Thus, the fitnesses of females (in the order above) are (1,1,1, 1+*s_f_*, 1+*s_f_*, 1+*s_f_*,) and the male fitness costs (in the order above) are (1,1-*s_m_*, 1, 1-*s_m_*+*s_x_*).

As in Model 1, whenever *s_f_* > 0 the *m* mitochondrial type invades when it is initially rare, and increases in frequency all the way to fixation, regardless of male fitnesses. In this simplified parameter space, whenever *s_x_ > s_m_*, the X-linked restorer will invade and go to fixation. The interesting contrast to Model 1 lies in the comparison of rates of increase of the Mother’s Curse mitochondria and of the restorer allele, and these questions are explored numerically below (Fig. 2). Allowing a more generalized assignment of fitnesses readily produces quite complex behavior, including protected polymorphism where neither the Mother’s Curse allele nor the restorer go to fixation but instead both cycle in frequency indefinitely (Rand et al. 2001).

#### Model 3 Mitochondrial Mother’s Curse and Y-linked restoration

The Y chromosome being strictly paternally inherited is a promising candidate location for restorers of male fitness in the face of the Mother’s Curse. However, the dynamics of this interaction has never as far as we are aware been formally modeled. Some empirical evidence suggests that mito-Y interactions may be an important determinant of male fitness. Comparing nuclear autosomal genes whose expression has been shown to be sensitive to either mitochondrial variation (Innocenti et al., 2011) or Y-linked variation (Lemos et al., 2008) reveals considerable overlap (Rogell et al., 2014). Direct tests of mito-Y interactions have come from experiments using crosses between mitochondrial and Y chromosome replacement lines in *Drosophila melanogaster* (Dean et al., 2015; Yee et al., 2015). These studies find extensive mito-Y epistasis, which at least partly appears to be environmentally sensitive, but to date there are no reports of the presence of Y-linked restorers of male fitness from the Mother’s Curse.

We consider a pair of mitochondrial alleles (*M* and *m*) and a pair of Y chromosomes (*Y* and *y*). Females are of two genotypes (*M-XX* and *m-XX*). Males are of four genotypes (*M-XY*, *m-XY*, *M-Xy*, and *m-Xy*). We imagine the mitochondrial mutation as having an advantage in females, so the fitnesses of females (*M* and *m*) are (1 and 1+*s_f_*) respectively. In males, the *m* mitochondrial allele comes at a fitness cost of *s_m_*, but only with the ancestral Y chromosome, and the *y* allele provides fitness compensation of *s_y_*. Thus the four male genotypes (in the order above) have respective fitnesses (1, 1-*s_m_*, 1 and 1-*s_m_*+*s_y_*). Because of the simple transmission rules of mtDNA and the Y chromosome, the recursion for the frequency of the *m* mitochondrial type (whose frequency is *p_m_*) is:

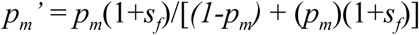

From the initial condition of *p_m_* = 0, the invasion condition for the novel Mother’s Curse mitochondria to have the eigenvalue of the linearized system exceed 1, and this occurs whenever *s_f_*> 0. Thus, whenever there is any advantage to the daughters, the new mitochondrial type invades and goes to fixation, regardless of its effect in males. The recursion for the male-fitness restoring *y* allele is:

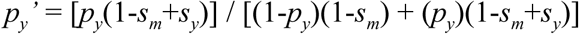

Here the fixation of *p_y_* = 0 is unstable, and both invasion and fixation of the restorer Y chromosomes is guaranteed when *s_y_ > s_m_*. There are no stable polymorphic states, and in fact this model has very rapid and simple dynamics, so the proportion of time the population is in transient polymorphism would be small. The fate of the population seems to reside in a mutation-limited state, where the waiting time to the next Mother’s Curse and next Y restorer will determine the balance of male and female fitnesses.

#### Model 4 Mitochondrial Mother’s Curse and Z-linked restoration

In a ZW system, the Z chromosome spends ⅔ of its time in males and only ⅓ of its time in females. As a consequence, the masculinized Z (Wright et al., 2012) may be expected to harbor male-biased restorers of Mother’s Curse (Wade and Drown, 2016). A recent study of the chromosomal location of nuclear genes with mitochondrial function, however, did not find any overrepresentation of such genes on the Z (Dean et al., 2014). Nevertheless, as this model shows, there remains good motivation to perform a screen for Z-linked variants that maybe have Mother’s Curse restorer function.

Following the nomenclature that is familiar by now, females will have four genotypes (*M-ZW*, *M-zW*, *m-ZW*, and *m-zW*). Males are of six genotypes (*M-ZZ*, *M-Zz*, *M-zz*, *m-ZZ*, *m-Zz*, and *m-zz*). The fitnesses of the females (in the order above) are (1, 1, 1+ *s_f_*, 1+*s_f_*) and the males are (1, 1, 1, 1-*s_m_*, 1-*s_m_*+*s_z_*/2, 1-*s_m_*+*s_z_*). Whenever *s_f_*> 0 the derived mitochondrial mutation *m* confers an advantage to females, and this comes at a cost to males of *s_m_*. If males acquire the derived *z* allele, their fitness is restored by *s_z_*/2 in *Zz* heterozygotes and by *s_z_* in *zz* homozygotes. The recursions for the allele frequency dynamics for both mitochondrial type that causes the female-favoring male-disfavoring effects and the Z-linked allele that restores male fitness are presented in the Supplement, along with the stability conditions of the fixation point where the frequencies of these alleles (*p_m_,p_z_*) are (0,0). Even in the absence of the (*Z,z*) polymorphism, the *m* mitochondrial type invades if *s_f_*>0, and the *z* allele accelerates fixation of the *m* allele (Fig. 2).

#### Model 5 Mitochondrial Mother’s Curse and W-linked restoration

The mitochondrial genome and the W chromosome are always co-transmitted and any mito-W combination that improves female fitness should spread. An explicit screen of mito-W interactions did detect some W-linked mito-nuclear genes in the blood fluke *Schistosoma mansoni*, but found no overrepresentation of these genes on the W (Dean et al., 2014). In addition, the low levels of mitochondrial genetic variation in birds compared to mammals has been suggested to be due to Hill-Robertson effects on the W (Berlin et al., 2007).

We consider a pair of mitochondrial alleles (*M* and *m*) and a pair of W-linked alleles (*W* and *w*). In this case, we assume no segregating variation on the Z. Females can be of four genotypes (*M-ZW*, *m-ZW*, *M-Zw*, and *m-Zw*). Males are two genotypes (*M-ZZ* and *m-ZZ*). Because the males always transmit the Z chromosome, and they do not transmit their mtDNA, the males essentially become irrelevant in the model, apart from assuming that even with an extremely biased sex ratio, all females can be fertilized. To be consistent with the XY Mother’s curse, the fitnesses of the *M-ZW* and *M-Zw* females are both 1, and the *m-ZW* and *m-Zw* females have fitness 1+ *s_f_*. The male fitnesses are (in the order above) 1 and 1 - *s_m_*.

This case behaves essentially like clonal evolution, as each female transmits the mitochondrion and W chromosome to all of her female offspring. Thus, trivially, whichever female genotype has the highest fitness always goes to fixation. The W chromosome and mitochondrion are inseparable with respect to sex-specific functional mutations that they may harbor, and each can drive the other’s selective sweeps. This observation raises the empirical question of whether there are W-mtDNA epistatic interactions, regardless of whether they cause or resolve sexual conflict.

### Father’s Curse

#### Model 6 Father’s Curse with Y variation driving a female deleterious autosomal allele

Here, we are introducing a novel kind of sexual conflict, which we call Father’s Curse. This curse arises when a derived Y-linked allele impacts expression of an autosomal locus such that one allele that favors males over females. A potential biological example is the high heterochromatin content of the Y chromosome of *Drosophila,* which has long been known to influence the chromatin state of the *white^mottled4^* allele and suppress position effect variegation (reviewed in Elgin and Reuter, 2013). Subsequently, it has also been demonstrated that differences in heterochromatin composition of the Y can impact gene expression and chromatin state of many regions of the genome (Lemos et al., 2010, 2008; Silkaitis and Lemos, 2014), consistent with the occurrence of Y chromosome driven sexual conflict.

Consider Y-linked alleles (*Y* and *y*) and autosomal alleles (*A* and *a*). Females are of only three genotypes (*AA*, *Aa* and *aa*), and males are of six genotypes (*AA-Y*, *Aa-Y*, *aa-Y*, *AA-y*, *Aa-y*, and *aa-y*). Female fitnesses are (1, 1-*s_f_*/2, and 1-*s_f_*) and male fitnesses are (1, 1, 1, 1, 1+*s_m_*/2, 1+*s_m_*). So the *y* allele has an advantage for *aa* males, that may be big enough to drive the autosome to an allele frequency that is disfavorable in females.

The dynamics of the Y chromosome are simple, owing to its haploid transmission. If the mean fitness of the bearers of the *y* allele exceeds the mean fitness of the bearers of the *Y* allele, then the *y* allele will increase in frequency. As the *y* allele increases in frequency, the shifting proportions of *y* and *Y* alleles may alter the marginal fitnesses of the autosomal genotypes, but given the constraints on the fitnesses, the *y* allele will always fix if it can invade. While on this trajectory toward fixation, the *y* allele can result in driving up the frequency of an autosomal allele in both sexes, even if that allele is disfavorable in females. Hence the name Father’s Curse.

One interesting behavior occurs when *s_f_* is sufficiently large compared to *s_m_*, and the autosomal allele is lost. The Y alleles may continue to segregate in the population, but they become effectively neutral. This underscores how Father’s Curse requires an autosomal locus that harbors the alleles with the sexual conflict, and under some circumstances the Y variant can serve to drive the male-favoring (and female-disfavoring) allele to fixation (Fig. 3). For further details of the dynamics of this model, see the Supplement.

**Figure 3.**
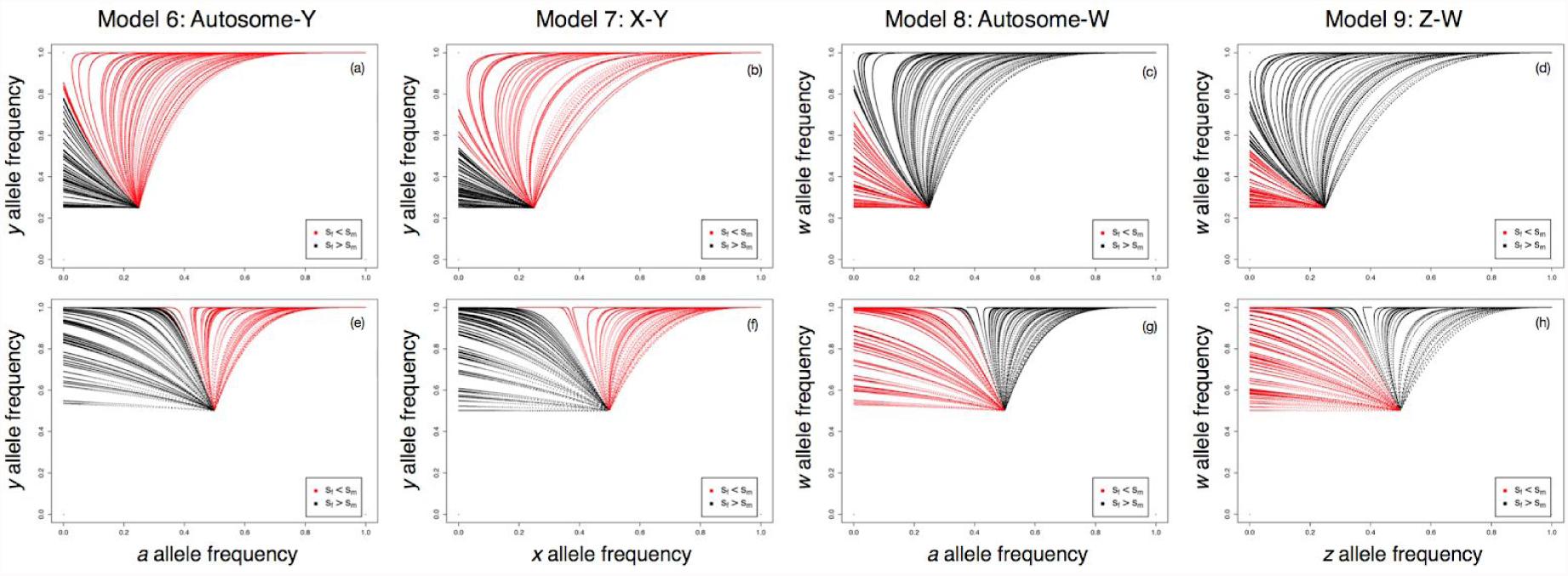
In Father’s Curse (Models 6 and 7) and Nuclear Mother’s Curse (Models 8 and 9), the uniparentally inherited sex chromosome (Y or W) can drive the fixation of an allele that is deleterious in the homogametic sex. Analytical work examines invasion of the sex-specific alles (Y and W) and the sexually antagonistic alleles (X, Z, or autosomal) from the lower left corner, but higher frequency starting points better illustrate the dynamical behavior. For Models 6-9, both alleles start with an initial frequency of either 0.25 (a-d) or 0.5 (e-h), and the change in allele frequency at every generation is tracked for a total of either 5000 (a,e,c,g) or 10,000 (b,f,d,h) generations. Simulations were run for 100 randomly selected combinations of *s_f_* and *s_m_* for each model. Trajectories where the absolute value of the selection coefficient in females (*s_f_*) is less than that in males (*s_m_*) are shown in red, and the opposite in black.

#### Model 7 Father’s Curse with Y variation driving a female deleterious X-linked allele

The effect of the Y chromosome on nuclear gene expression is not restricted to autosomes. Experimental work in *D. melanogaster* has shown that Y-linked genes can alter the expression of genes on the X chromosome (Jiang et al., 2010). In these studies, genes on the X that were located close to the euchromatin–heterochromatin boundary were particularly sensitive. Here, we consider the case in which a novel variant of the Y chromosome impacts expression of an allele at an X-linked locus that favors males over females. Co-evolution of the X and the Y results in increase in frequency of the male-favoring X at the cost of displacing females from their fitness optimum.

Consider two Y alleles (*Y* and *y*) and two X-linked alleles (*X* and *x*). Females are of only three genotypes (*XX*, *Xx* and *xx*), and males are of four genotypes (*XY, Xy, xY,* and *xy*). Females fitnesses are (1, 1-*s_f_*/2, and 1-*s_f_*) and male fitnesses are (1, 1, 1, 1+*s_m_*). So the*y* allele has an advantage for *xy* males, that may be big enough to drive the X-locus to an allele frequency that is disfavorable in females. Together with the autosome-Y example described above (Model 5) this provides another examples of how a father’s alleles serve to favor sons at the expense of the daughter’s fitness, and thus of the phenomenon we dub Father’s Curse.

The Supplement establishes the recursions for this system, which we linearize to determine the conditions for invasion of the *y* allele. Because the initial Y chromosome allele dynamics depend only on the males, if *s_m_* > 1, the *y* allele can invade, and this is consistent with the numerical results. The Y chromosome allele dynamics are very rapid because of its haploid-like transmission, and as it invades it drags the female-deleterious *x* allele with it (Fig. 3).

### Nuclear Mother’s Curse

#### Model 8 Nuclear Mother’s Curse with W variation driving a male deleterious autosomal allele

Analogously to the Model 6, where the Y chromosome-autosome epistasis causes the spread of a female-harming autosomal mutation, the W chromosome may drive the invasion of a male-harming autosomal mutation. This is a form of Mother’s Curse, but unlike previously described examples, it is not due to a cytoplasmic (mitochondrial) mutation. We call this a nuclear Mother’s Curse. We have yet to find a study that sought to detect this effect in a natural population, and such a study would appear to be well motivated.

Consider two W alleles (*W* and *w*) and autosomal alleles (*A* and *a*). Males have three genotypes (*AA*, *Aa*, and *aa*), females are of six genotypes (*AA-W*, *Aa-W*, *aa-W*, *AA-w*, *Aa-w*, and *aa-w*). Male fitnesses are (1, 1- *s_m_* /2, and 1- *s_m_*) and female fitnesses are (1, 1, 1, 1, 1+ *s_f_*/2, 1+ *s_f_*). The *w* allele provides and advantage for the *Aa* and *aa* females, which may be big enough to make a spread despite its fitness cost in males. Contrasting this model with Model 6 (Y-autosome Father’s Curse) is instructive. Examination of the fitnesses in Table 1 reveals that the two models are equivalent, except for the sex labels, and this is consistent with the recursions and the appearance of the simulation results in Figure 3. Of course, biologically these two situation may not be expected to be so analogous, given the large difference in the variance in offspring numbers of males and females, the expected greater impact of sexual selection in males than females in most species, and the unknown degree to which the W chromosome impacts chromatin state and gene expression genome-wide.

#### Model 9: Nuclear Mother’s Curse with W variation driving a male deleterious Z-linked allele

In this model, variation on the W chromosome interacts with a Z-linked locus to favor alleles that are female-advantageous and male disadvantageous to cause a second kind of nuclear Mother’s Curse.

Consider W alleles (*W* and *w*) and Z-linked alleles (*Z* and *z*). Males are of only three genotypes (*ZZ*, *Zz*, and *zz*). Females are of four genotypes (*ZW*, *zW*, *Zw*, and *zw*). Males fitnesses are 1, 1-*s_m_*/2, and 1-*s_m_* and female fitnesses are 1, 1, 1, and 1+*s_f_* respectively. Inspection of these fitness parameters shows that it is the sex-flipped version of Model 7 (X-Y Father’s curse), with the same parameterization. As mentioned above for the contrast of Models 6 and 8, aspects of the biology will likely prevent Models 7 and 9 from being biologically equivalent, but with respect to the models presented here, the stability conditions and dynamics will be the same (see Fig. 3). This means that the dynamics of the W chromosome will be more rapid than the Z, and the conditions for the initial invasion of the *w* allele depend on polymorphism at the Z locus. Further details can be found in the Supplement.

## Discussion

In their highly cited letter to *Nature*,Frank and Hurst (1996) describe the logic of the population genetics argument for what we now call Mother’s Curse as “indisputable.” In this paper, we have extended this logic to consider the full spectrum of paternal curses caused by the uniparental inheritance of genes. Using a combination of analytical models and computer simulations, we determine the fitness consequences of transmission asymmetries of nuclear, mitochondrial, and sex chromosome-linked genes and how they may both cause and resolve sexual conflicts. Contrary to recent arguments the evolution of nuclear restorer genes is an unlikely route to reverse the mother’s curse (e.g. Wade, 2014) we show how restorers can readily evolve, but the rate depends on the exact transmission pattern of the nuclear chromosome where the restorer is located. We also demonstrate a novel way by which interactions between a uniparentally and biparentally inherited genes can lead to sexual conflict: Father’s Curse and nuclear Mother’s Curse.

The general framework developed here opens up many avenues for informative extensions. In a recent paper,Connallon et al. (2018) developed a quantitative measure of male mitochondrial load–the reduction in fitness due to the accumulation of mitochondrial mutations–in the context of Mother’s Curse. This effort is well motivated by the observation that the dynamics of the model are relatively simple, and so an important feature is the steady state frequency of curse alleles in transition to fixation or loss. It also motivates a contrast of the levels of load across the three classes of parental curse models outlined here, including mitochondrial Mother’s Curse, nuclear Mother’s Curse, and Father’s Curse.

Demographic processes may alter some of the trajectories described here. Our models assume random mating, but Wade and Brandvain (2009) and Unckless and Herren (2009) have both showed how inbreeding, mating between close relatives, will limit the spread of mitochondrial Mother’s Curse variants because it links a mother’s fitness with that of her sons. Similarly, if males affect the fitness of their sisters, kin selection may have the same effect (Wade and Brandvain, 2009). In addition to altering the trajectory of the Mother’s Curse allele, we also expect the spread of male restorers to be affected by these processes.

In all our models we have assumed that male gametes are never limiting, which is formally equivalent to assuming that the sex ratio remains 50:50 throughout. Hamilton (1967) showed how a driving Y chromosome can result in a male biased (“extraordinary”) sex ratio, which may eventually drive the population extinct. Since then, several sex chromosome drivers have been identified (reviewed in e.g. Bravo Núñez (2018)). Maternally inherited mitochondrial mutations should yield a female-biased sex ratio, which has also been suggested as an alternative explanation to Haldane’s Rule (Hurst and Pomiankowski, 1991). A biased sex ratio will likely affect the dynamics of our models, and relaxing the 50:50 assumption would allow us to incorporate another kind of genetic conflict.

In Models 1-4 we are modeling the fate of a nuclear mutation restoring male fitness and we assume the restoration is associated with no cost. However, this assumption is questionable. The cost of nuclear restorers has been best explored in the context of the evolution of gynodioecy, the presence of females and hermaphrodites in a population (Bailey and Delph, 2007; Delph et al., 2007). Studies in a variety of plant systems have characterized the molecular mechanism and fitness cost of nuclear restorers of male fertility in hermaphrodites. A recent example is the study of the cytoplasmic male sterility system in *Brassica napus* by Montgomery et al. (2014), which demonstrated not only the presence of costs but also showed how the fitness cost differed between two different nuclear restorers. Without such costs, theory predicts that restorers will always fix and they are therefore crucial to the maintenance of gynodioecy (Delph et al., 2007). As we learn more about the fitness consequences of mitochondrial mutations, including in animals, and how they depend on the nuclear background, incorporating the cost of nuclear restorers could be a productive focus of future modeling efforts.

As is typical for these kinds of models, we assume no paternal leakage (inheritance from the father) of the mitochondrial genome. While maternal inheritance of the mitochondrial genome is shared across the tree of life, exceptions exist. Molluscan bivalves, for example, have doubly uniparental inheritance, whereby females pass on their mitochondrial genomes to both sons and daughters, whereas males only transmit to sons (Breton et al., 2007). In some plants, including cucumber (Havey, 1997) and many conifers (Worth et al., 2014), uniparental paternal inheritance appears to be the norm. Lastly, paternal leakage has also been reported in variety of systems, including but not limited to humans (Schwartz and Vissing, 2002), fruit flies (Wolff et al., 2013), sheep (Zhao et al., 2004), and chicken (Alexander et al., 2015). Paternal leakage will introduce direct purifying selection on the male deleterious mitochondrial mutations, therefore limiting the spread such mutations. Paternal leakage, however, appears to be rare (an early estimate in mice put it at 10^−4^; (Gyllensten et al., 1991)) and it is generally considered too weak of a force to reverse the Mother’s Curse (Connallon et al., 2018; Engelstädter and Charlat, 2006; Wade and Brandvain, 2009).

Our models also have implications for the potential role of transmission asymmetries in generating reproductive isolation through Bateson-Dobzhansky-Muller incompatibilities. The Bateson-Dobzhansky-Muller model (Bateson, 1909; Dobzhansky, 1936; Muller, 1942) characterizes the emergence of incompatibilities between two allopatric populations due to the fixation of variants within one population that, while neutral in their original background, are deleterious in the genetic background of the other. Burton and Barreto (2012) highlight numerous such incompatibilities between mitochondrial and nuclear genomes that result in hybrid breakdown, and the evolution of pairwise incompatibilities among mitochondrial, autosomal, and X-linked genes in parapatric populations has also been further explored through analytical models (Höllinger and Hermisson, 2017). While our models only consider a single population, the potential rapid fixation of both alleles in every model highlights the potential accumulation of two variants that in combination are either neutral or advantageous.

Mating between two previously isolated populations could disrupt these interactions leading to hybrid incompatibilities. Extensions of our models that incorporate multiple populations with migration among them may therefore provide further insight into how asymmetrically inherited genomic components contribute to the genetics of reproductive isolation.

## Acknowledgements

This work was supported by a Meinig Family Professorship for AGC, donated by Nancy and Peter Meinig to Cornell University. JAÅ was supported by a fellowship from the Sweden-America Foundation.

## Supplement

Here we consider the stability conditions for the ancestral state in which the population has neither the sexual conflict allele (causing either Mother’s Curse or Father’s Curse) nor the restorer allele.

### Model 4 Mitochondrial Mother’s Curse with a Z-linked restorer

For this model the fitnesses are:

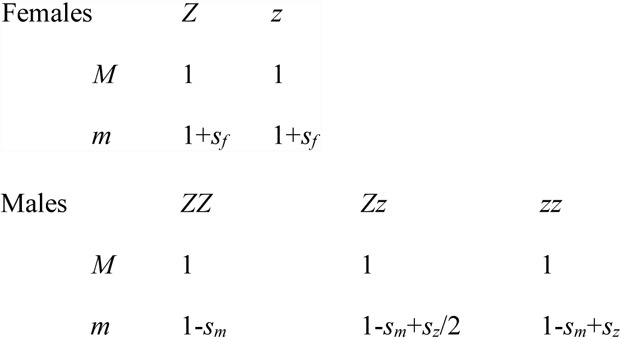

so the *m* mitochondrial type has advantage *s_f_* for females and disadvantage *s_m_* for males (assuming *s_f_, s_m_* > 0). The *z* allele has no effect in females but restores male fitness by an amount *s_z_*/2 in heterozygotes and sz in *zz* homozygotes. Letting the frequencies of the *m* and *z* alleles be *p_m_* and *p_z_*, we see that near the point (*p_m_*=0, *p_z_*=0) the linkage-disequilibrium-like term describing the association between *p_m_* and *p_z_* is negligible, and descriptions of initial frequency dynamics can ignore this term. As a consequence, we can determine the stability of this point from the recursions in the two allele frequencies:

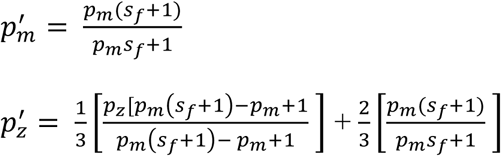

To examine the stability characteristics of this model, at the point (*p_m_* = 0, *p_z_* = 0), we determine the leading eigenvalue of the Jacobian matrix:

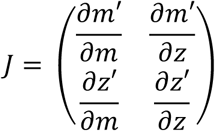

Evaluating the Jacobian at (*p_m_* = 0, *p_z_*=0) yields a leading eigenvalue of *s_f_*+1, showing that whenever *s_f_* > 0, this corner equilibrium is unstable and the Mother’s Curse mitochondrial type will invade. When the *m* mitochondrial type gets to fixation, which it will do relatively quickly, the *z* allele behavior can be determined by examining the (*p_m_*= 1, *p_z_*=0) corner equilibrium. This is unstable whenever *s_z_*>0, implying that the restorer allele will invade. The actual dynamics in the middle of this phase space is a bit more complex, and can be seen in Figure 2 of the main text.

### Model 6 Father’s Curse with Y allele driving an autosomal female-deleterious allele

For this model the fitnesses are:

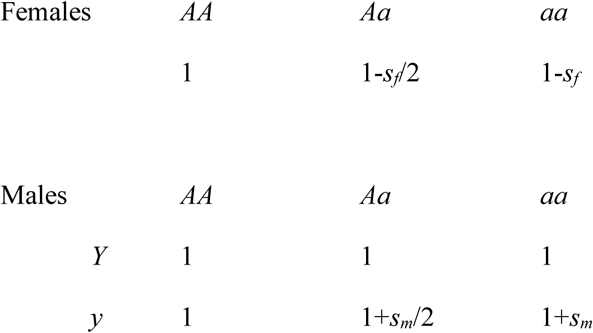

The recursion for the autosomal *a* allele is:

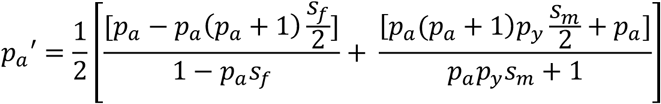

The recursion for the *y* allele is:

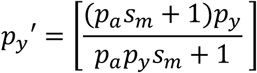

Evaluating the Jacobian at *p_a_* = 0, yields a leading eigenvalue of (*p_y_s_m_* – *s_f_* +4)/4, showing that a female advantageous allele can invade if *s_f_* < 0 as *p_y_* → 0.

### Model 7 Father’s Curse with Y allele driving an X-linked female-deleterious allele

For this model the fitnesses are:

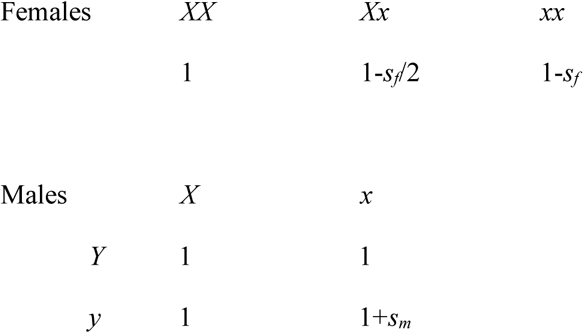

Here the recursion for the X-linked *x* allele is:

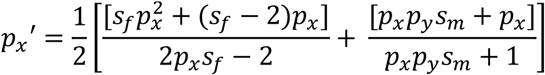

And the recursion for the Y-linked *y* allele is:

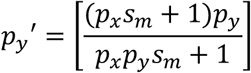

Stability analysis proceeds as above, solving the leading eigenvalue of the linearized recursion. Provided the *p_x_* > 0, this eigenvalue is simply 1+*s_m_*, implying that the Father’s Curse allele can invade when there is a fitness advantage to the sons, even at the fitness expense *s_f_* to the daughters. While the increase in frequency of *y* is monotonic, if *s_f_* is sufficiently large, the *x* allele is lost, and at this point, the Father’s Curse *y* allele becomes effectively neutral and it stops increasing in frequency. This can also be seen in the phase diagram of Figure 3.

### Model 8 W-autosomal nuclear Mother’s Curse

Here the female’s fitness is improved by a novel allele (*w*) of the strictly female-transmitted W chromosome, and the *a* allele at an autosomal locus restores male fitness by *s_m_*/2 in *Aa* heterozygotes and by amount *s_m_* in *aa* homozygotes. The full model has these fitnesses:

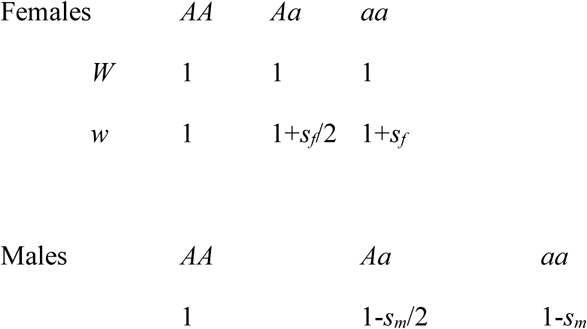

Here the recursion for the *w* allele frequency is entirely determined by females, and is:

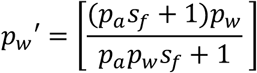

And the recursion for the autosomal restorer allele is:

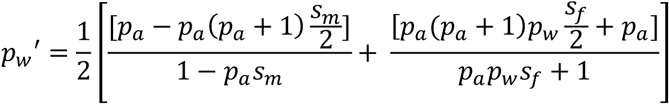

Stability conditions have already been established for Model 6, and here we have the *w* allele invading only if it can drive a sexually antagonistic autosomal allele (female-favoring and male-disfavoring). Whenever the *w* allele goes to fixation, the *a* allele is also fixed, although allele frequencies need not change monotonically. If the *a* allele is lost, the fitnesses collapse to all being 1, and further selection on the *w* allele is arrested (and its dynamics will be neutral).

### Model 9 W-Z nuclear Mother’s Curse

Here, as in Model 8, the female’s fitness is improved by a novel allele (*w*) of the strictly female-transmitted W chromosome, and the *z* allele at a Z-linked locus restores male fitness by *s_m_*/2 in *Zz* heterozygotes and by amount *s_m_* in *zz* homozygotes. As mentioned above, the model is formally identical to Model 7, replacing the Y locus with the W locus, and flipping the sex labels. The full model has these fitnesses:

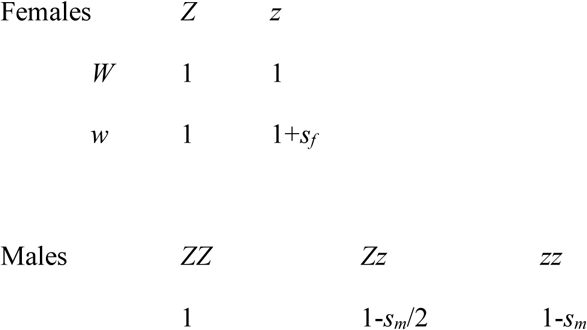

Here the recursion for the Z-linked *z* allele is:

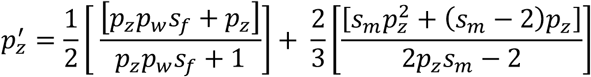

And the recursion for the W-linked *w* allele is:

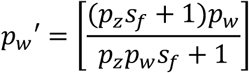

The dynamics and conditions for invasion of the nuclear Mother’s Curse are identical to the conditions for invasion of Model 7, the Father’s Curse with an X-linked locus having the sexual conflict. Here, the *w* allele can fail to fix if the z allele is first lost.

